# Structure-Based Discovery of a NPFF1R Antagonist with Analgesic Activity

**DOI:** 10.1101/2023.10.25.564029

**Authors:** Brian J. Bender, Julie E. Pickett, Joao Braz, Hye Jin Kang, Stefan Gahbauer, Karnika Bhardwaj, Sian Rodriguez-Rosado, Yongfeng Liu, Manish Jain, Allan I. Basbaum, Bryan L. Roth, Brian K. Shoichet

**Affiliations:** Department of Pharmaceutical Chemistry, University of California, San Francisco, CA 94158, USA; Department of Pharmacology, University of North Carolina School of Medicine, Chapel Hill, NC 27599, USA; Department of Anatomy, University of California, San Francisco, CA 94158, USA

**Keywords:** NPFF1R, structure-based drug discovery, small molecule, analgesia

## Abstract

While opioid drugs remain among the most effective analgesics for pain management, adverse effects limit their use. Molecules that synergize with opioids, increasing analgesia without increasing side effects, could prove beneficial. A potential way to do so is via the RF-amide receptor system, as NPFFR1 agonists reduce µ- opioid receptor (µOR)-based analgesia while antagonists increase it. These inferences are, however, clouded by the lack of selectivity of most NPFF1R ligands. Seeking selective antagonists of the NPFF1R, we screened a large virtual library against a homology model of NPFF1R. From 26 high-ranking molecules that were synthesized and tested, one antagonized NPFF1R with a K_i_ of 319 nM. Structure-based optimization led to a 22 nM antagonist of NPFF1R, compound **56**, with selectivity against a large panel of GPCRs. When administered alone, **56** has no activity in mouse tail-flick nociception assays. However, coadministration of compound 56 and morphine produced significantly greater antinociception than did morphine alone, consistent with the notion that NPFF1R nociceptive activity occurs via modulation of µOR signaling. Surprisingly, in the hot-plate assays **56** was analgesic by itself, suggesting that NPFF1R alone can also confer analgesia. At equi-analgesic doses, combinations of **56** with morphine reduced the common constipation side effect of morphine versus using morphine alone. The high selectivity of **56** and its activity in cooperation with morphine supports further analgesic development against NPFF1R and against the RF-amide family of receptors more generally.

## Introduction

The RF-amide receptors are a family of five G protein-coupled receptors (GPCRs) that are activated by endogenous peptide hormones harboring a C-terminal RF-amide, or Arg-Phe-NH_2_, motif^1^. These peptides and their corresponding receptors are conserved through evolution and are implicated in cardiovascular functions, energy homeostasis, and feeding regulation^2–4^. Soon after their initial discovery in mammalian lineages, their contribution to pain processing was reported. Injection of mice with the peptides NPFF or NPAF is pro-nociceptive, decreasing latencies in a tail-flick assay^5^ and NPFF injection also reduced the analgesic effect of morphine. This anti-analgesic action could be reversed by administration of BIBP-3226, a non-selective NPFF1R and NPFF2R antagonist or by an Y1-neuropeptide Y receptor type 1 (NPY1R) antagonist^6^. Administration of BIBP-3226 enhanced the analgesia produced by morphine, and indeed cyclic peptides have been designed to modulate both opioid and NPFF receptors simultaneously ^7^. Taken together those results suggested that antagonism of the RFamide receptors NPFF1R or NPFF2R may be useful in the treatment of pain via co-activity with µ-opioid receptor signaling^8,9^ and that there may be an ongoing pronociceptive action at RF-amide receptors.

To understand the contribution of the RF-amide receptors to nociception, selective agonists and antagonists are required. Whereas both peptides like NPFF and peptidomimetic antagonists like BIBP-3226^6,10^, RF-9^8^ and newer antagonists like compounds **22e** and **12**^11^ have been very useful, each has potential drawbacks as molecular probes. Even the native peptide agonists lack selectivity within the family with, for instance, NPFF activating both NPFF1R and NPFF2R. Furthermore, the antagonist BIBP-3226 is large (MW 474 amu) with a mixed cationic and amphiphilic character, limiting its pharmacological uses, and it not only antagonizes NPFF1R, but also NPFF2R and NPY1R with similar potencies as it was developed for the NPY1 receptor, which is itself a pain target ^12^. The antagonist RF-9 has similar liabilities. While the sub-nanomolar antagonist compound **12** is highly selective for NPFF1R, it inherits the high molecular weight (500 amu), hydrophobicity (cLogP of 3.2), and dipeptide nature of several of the other series. The antagonist **22e** has shed this peptidic nature, but it has similar mid-nanomolar antagonism of both NPFF1R and NPFF2R^11^ We thus thought it would be interesting to explore NPFF1R’s role in augmenting opioid analgesia with selective NPFF1R antagonists with favorable physical properties and pharmacokinetic exposure.

To do so, we adopted a structure-based approach, docking a library of 166 million make-on-demand or “tangible” molecules against a three-dimensional model of the receptor. By insisting on fragment-and lead-like molecules^11 13^ (molecular weight ≤ 350 amu, cLoP ≤ 3.5) we ensured that candidates would have favorable physical properties. A challenge with NPFF1R was that we did not have an experimental structure against which to dock, and so we created a homology model. Docking into homology models introduces errors relative to experimental structures, and these may be compounded by the challenges of a peptide receptor site and by the few known antagonists to use in control docking calculations, on which we typically depend^14^. Thus, though large library docking has found nanomolar and even sub-nanomolar ligands against the structures of hormone and transmitter-like receptors and enzymes^15–21^ hit rates and affinities have suffered against homology models of the same^22^ and, independently, against experimental structures of peptide receptors^23^. They have also been low versus modeled structures of orphan receptors with few control molecules^24,25^.

To mitigate these challenges, we created 1000 models of NPFF1R using Rosetta (see Methods). The model best suited for the large library docking screen was selected for its ability to prioritize the few known antagonists versus a larger set of property-matched decoys. Although these decoys resemble the known antagonists physically, they are topologically unrelated, and so unlikely to bind^14,26^. This led to mixed success; the hit-rate was lower than what the field has seen against the experimental structures of small-molecule receptors. Still, the antagonist that emerged was relatively potent and of low molecular weight compared to the known antagonists, with preliminary indications of selectivity. Optimization led to a 22 nM NPFF1R antagonist with selectivity against a panel of GPCRs. This antagonist was remained a fragment (molecular weight 215, cLogP 3.1) whose favorable physical properties ensured that it had good pharmacokinetic exposure on systemic dosing. This, combined with its selectivity, allowed us to investigate the role of NPFF1R in modulating the analgesia conferred via the µOR, in both wild-type (WT) and µOR knock-out mice. The usefulness of this antagonist as a selective NPFF1R probe, and prospects for further lead discovery against this target, will be considered.

## Results

### Homology model

To generate a suitable homology model, we used three templates with greater than 30% sequence identity to NPFF1R: Y1R, OX1R, and OX2R. To ensure the binding pocket remained open during energy minimization, the antagonist BIBP-3226 was co-modeled in the orthosteric site. The initial pose of BIBP-3226 was obtained from overlapping atoms in the ligand UR-MK-299 that was co-determined with Y1R (SF 1). One thousand models were calculated using Rosetta (see Methods); these models had relatively conserved transmembrane bundles, differing mostly in loops and termini (**Figure 1**). Each model was challenged to identify known actives from a pool of property-matched decoys^26,27^ as previously described^24,25^. Several models prioritized actives versus decoys, as measured by the logAUC enrichment, with the docked poses of the actives adopting reasonable poses. The models were further optimized for enrichment through modification of the dielectric boundary in the DOCK3.7 scoring grids (SF1)^12 14^. A final set of well-performing models was selected for a screen with the known ligands embedded in a library of about 1 million monocations selected from the larger library^14,27^. The model that best enriched the known ligands against this larger set was selected for the prospective large library docking.

**Figure 1.**
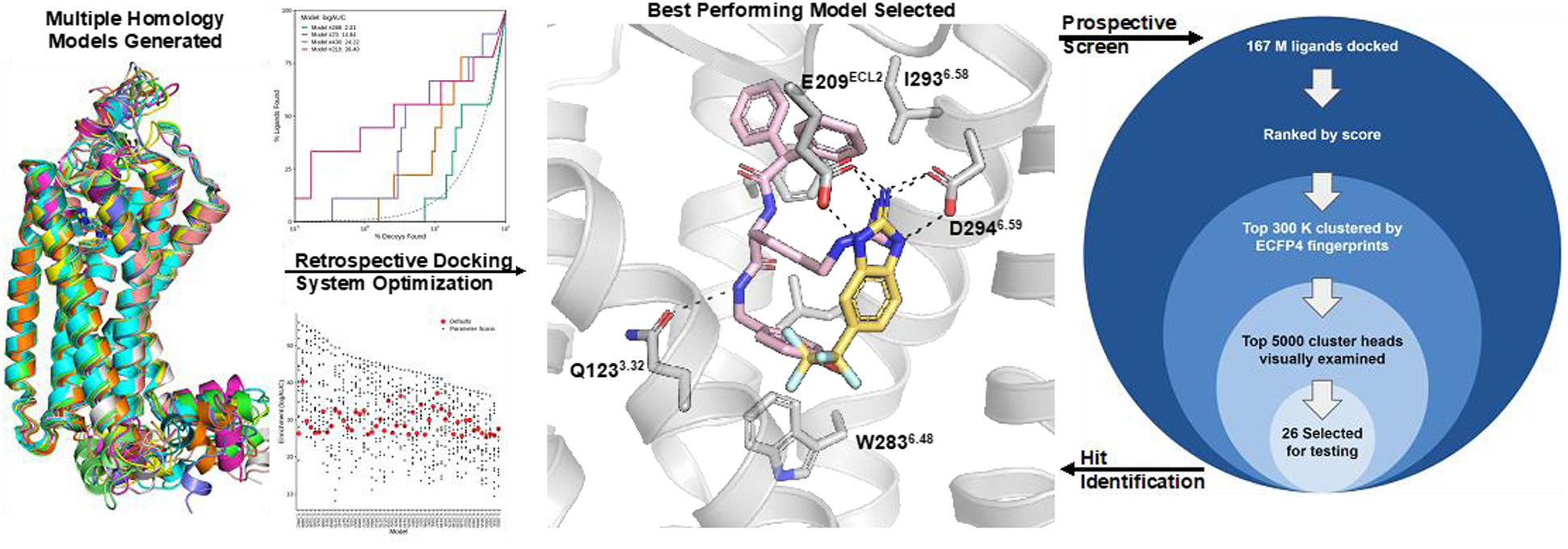
Docking workflow against NPFF1R. 1000 homology models were calculated for NPFF1R using the structure of NPY1R as a template. The model most suitable for docking was selected based on enriching known ligands against property-matched decoys, in sensible geometries. Against that model, 167 million molecules were docked and prioritized by score, diversity, and topological dissimilarity from the known ligands. The docked structure of the initial active, compound **16**, is shown in the middle panel (carbons in yellow) superposed on the structure of BIBP-3226. Key hydrogen bonding interactions are shown as dashed lines.

### Prospective large library docking

Based on the importance of the conserved arginine in RF-amide peptide agonists, and the cationic nature of several early RF-amide peptidomimetics like BIBP-3226 and RF-9^6,8^, in the docking campaign we focused on molecules modeled to be positively charged at physiological pH. A set of 166 million monocations from the ZINC20 library^26 28^ were docked into the best performing NPFF1R homology model. The top-ranking 300,000 compounds (top 0.18%) were clustered into sets with Tanimoto coefficients > 0.35 using ECFP4 fingerprints, and the best scoring compound from each cluster was identified. These cluster heads were ranked by DOCK score and the top 5000 were examined visually in the context of the binding pocket, using Chimera^29^. Compounds were prioritized if they occupied similar space and made similar interactions as the modeled BIBP-3226. In particular, ligands were prioritized if they formed a salt bridge with Asp294^6.59^, a conserved residue in RF-and RY-amide receptors. Conversely, compounds were deprioritized if they adopted strained torsion angles or high-energy tautomers. A set of 26 compounds, as far as we know not previously made, were synthesized from the virtual library and tested in signaling assays.

### Identification of ZINC725343470 as a 0.3 µM antagonist

The 26 compounds (**SI Figure 2**) were first tested at NPFF1R in a Tango functional assay^28 30^. Although most did not measurably antagonize the activity of the agonist peptide NPFF, one compound, **16** (ZINC725343470) did, with a K_i_ of 2.4 µM (**Figure 2**). Compound **16,** a 2- aminobenzimidazole with a pentafluoroethyl at the 5 position, is docked to ion pair with Asp294^6.59^ and Glu209 from ECL2, while its hydrophobic pentafluoro group is docked to form apolar interactions with Gly124^3.33^, Val127^3.3^ and Met173^4.57^ (**Figure 1**). Its small size ensures a high ligand efficiency of 0.54 kcal/HAC. This compound was confirmed in BRET assays between labeled receptor and miniGi with a Ki value of 0.3 µM. In dynamic light scattering (DLS) and in enzyme counter-screens (**SI Figure 3**), the compound does not aggregate at relevant concentrations, consistent with specific on-target activity at NPFF1R. We note that a 4% hit-rate is low by the standards of large library docking to hormone-and transmitter-binding GPCRs, where hit rates have often ranged from 17 to over 50%^15,16,18–21^. This lower hit rate likely reflects the homology model, the few good ligands to use in the docking control calculations, and the intentional diversity of the hit list—compound **16** was topologically unrelated to the rest of the initial docking hits; it had no analogs among the molecules prioritized for testing.

**Figure 2.**
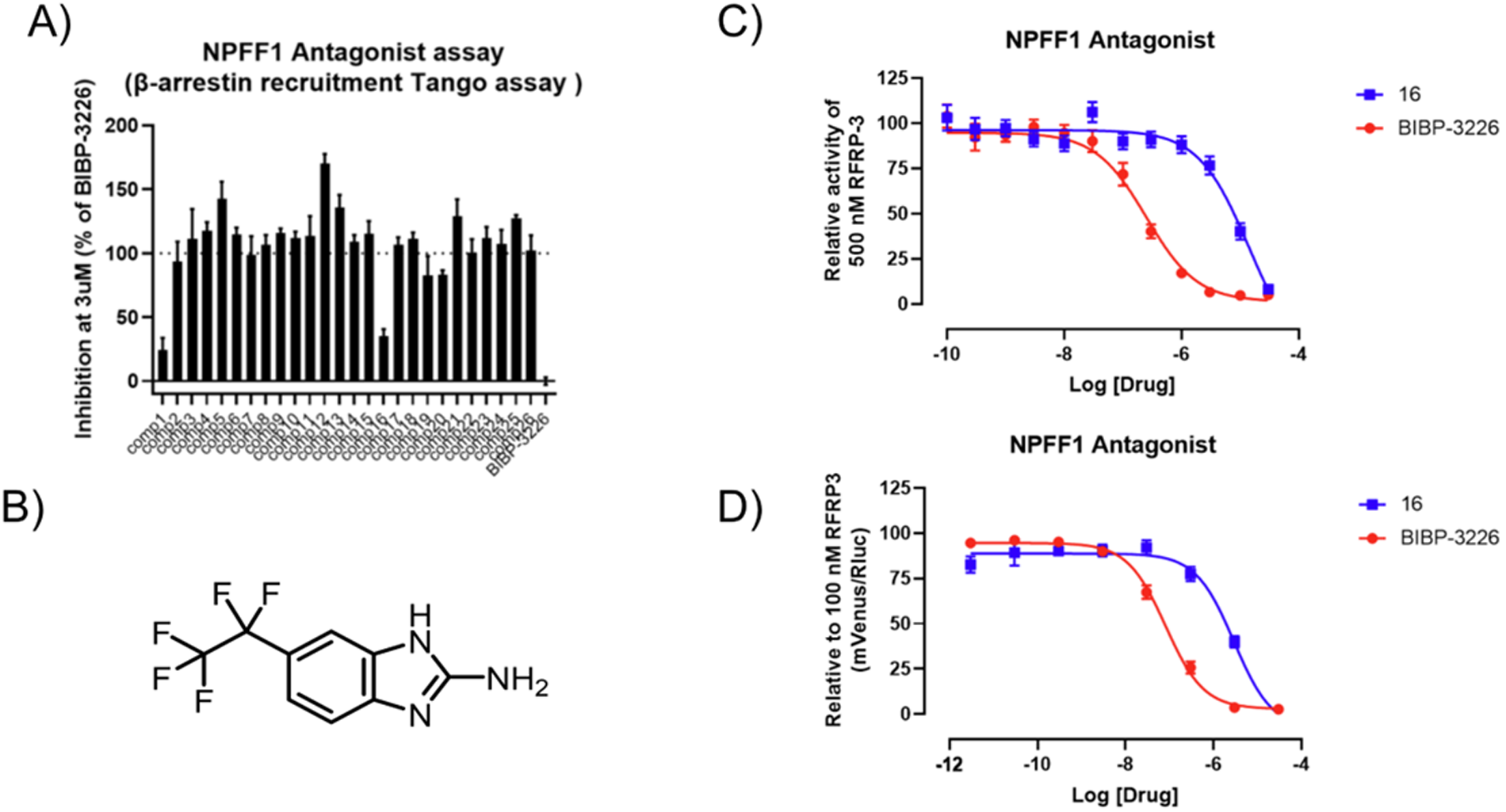
Screening Compounds at NPFF1R. (A) The 26 compounds from the virtual screen were tested in single-point inhibition of signal in the Tango assay at 3 uM. (B) The 2D structure of compound **16**. It was retested in full concentration response curve for Tango (C) and BRET (D) with Ki’s measured at 2.4 µM and 0.3 µM, respectively.

**Figure 3.**
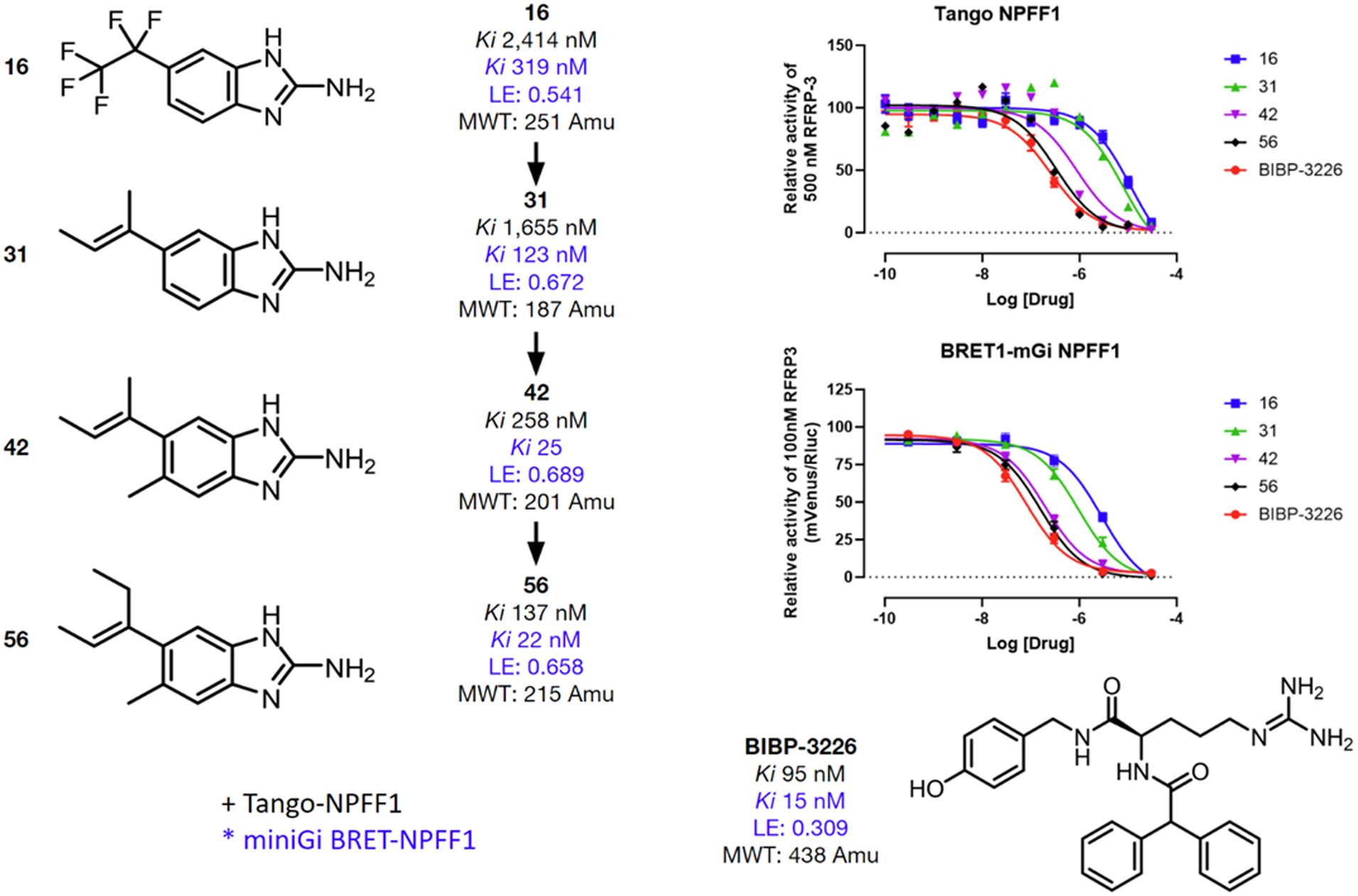
Optimization of compound 16. 2D structures of the progression from compounds **16**, **31**, **42**, and **56**, with corresponding potencies at Tango and BRET assays and ligand efficiency at BRET. Reference antagonist BIBP-3226 is also shown.

### Structure-based optimization

Based on its docked structure in the modeled site of NPFF1R (**Figure 1**), we sought to optimize **16**’s affinity while reducing its hydrophobicity. While **16** is potent for its size, its heavy fluorination is a liability. We thus first focused on modifying the 5-pentafluoroethyl, which would afford greater synthetic freedom and lower hydrophobicity. An analog that replaced the pentafluoroethyl with a simple ethylene, **29**, retained substantial activity in the BRET assay (**SI Table 1**), supporting the idea that this side chain could be modified. With the freedom to operate that this simple side chain afforded, we could then test the importance of the putative ion pair between the 2-amino-benzimidazole of the antagonist with Asp294^6.59^ and with Glu209^ECL2^. Replacing the ethylene of **29** with an ethyl, and methylating the terminal amine led to **28**, which lost all measurable activity. The same is true of a related analog that replaced the amino group entirely with a methyl (**27**). Both modifications disrupt the ionic interactions of the antagonist with the modeled Asp/Glu pair in NPFF1R. With the apparent essentiality of this ion pair supported, we returned to alkyl tail modifications.

The modeled structure of the NPFF1R/**16** complex suggested that a range of side chains could be tolerated at the 5 position, and potentially the 6 position. Ultimately 25 analogs were explored (**SI Table 1**). Based on the results obtained with **29**, the pentafluoroethyl of **16** was replaced by a pseudo-isosteric 2,3-butene (**31**). This eliminated any fluorophobic effect without sacrificing affinity, leading to a 123 nM antagonist (**Figure 3**). Replacing the 5-alkyl sidechain with amide sidechains (**37**, **38**, **40**) led to compounds without measurable activity, suggesting that while fluorophobic groups are not necessary, polar groups in this region of the binding site are unfavorable—this is sensible given the hydrophobic residues that are modeled to surround the antagonist in this region (e.g., Gly124^3.33^, Val127^3.36^, and Met173^4.57^). Adding a 6-methyl (compound **42**) improved Ki to 25 nM. Further addition of a single methyl to create a pentenyl-derivative yielded compound **56** with a Ki of 22 nM.

In our hands, BIBP-3226 and **56** inhibited NPFF1 similarly in the Mini-Gi BRET assay (**Figure 3**). As BIBP is known to antagonize NPY1R, we further screened these compounds at this receptor to measure off-target activity (**SI Figure 4**). BIBP-3226 was a full antagonist in this assay, as reported previously, while we found that **56** has no activity at NPY1R. Compound **56** was further counter-screened against a panel of 320 GPCRs at 10 µM and found to have only modest agonist activity at NPY5R (**SI Figure 4**); against all other targets in the panel, no significant agonist or inverse agonist activity was observed. Thus, compound **56** is a 22 nM antagonist of NPFF1R with little measurable agonism against a panel of 320 GPCRs. The molecule’s status as a fragment (molecular weight 215 amu) and its favorable physical properties (ligand efficiency of 0.66) made it a good candidate to advance *in vivo*, where its selectivity could help probe for the function of NPFF1R without confounds from its most similar off-targets.

**Figure 4.**
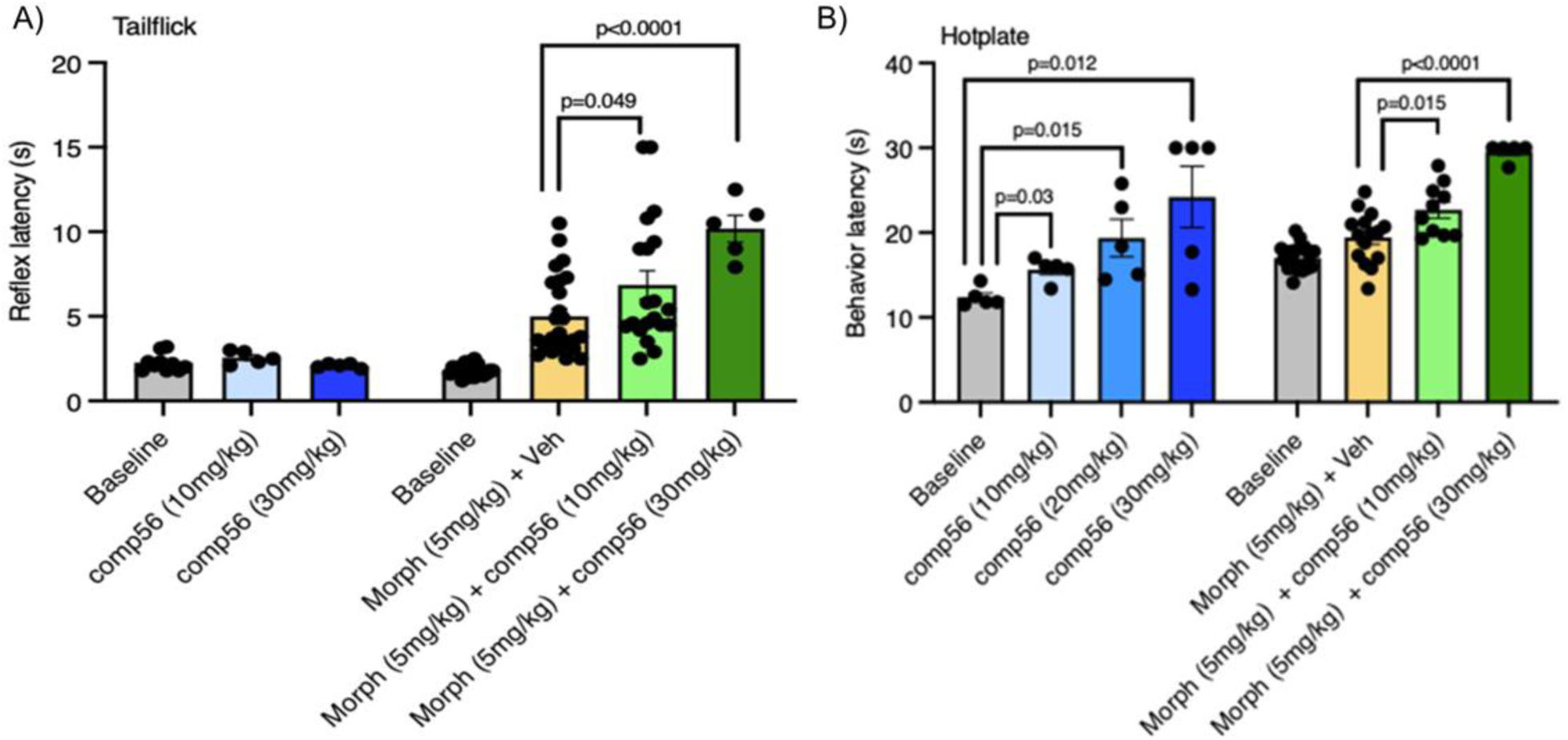
Antinociceptive effects of compound 56. (**A**) In the tail flick assay, 10 mg/kg (n=5) or 30 mg/kg (n=5) of i.p. compound **56** did not increase reflex latencies in the absence of morphine, but did so in combination with morphine. At constant i.p. morphine (5 mg/kg, n=25), latencies increased in a dose-dependent manner (10 mg/kg: n=20, 30 mg/kg: n=5) with increased compound **56** (i.p). (**B**) In the hot plate assay, i.p. injections of compound **56** increased latencies to paw withdrawal (analgesia) both alone (**left bars**, 10 mg/kg: n=5, 20 mg/kg: n=5, 30 mg/kg: n=5) and in combination with i.p. morphine (5 mg/kg, n=15 (**right bars,** 10 mg/kg: n=10, 30 mg/kg: n=5). Increasing doses conferred increasing analgesia. Data is shown as mean ± SEM. *p*-values between indicated groups were calculated with two-tailed unpaired *t*-tests.

### Pharmacokinetics

As a first step, we wanted to measure the exposure of compound **56** in the periphery and especially in the CNS, where anti-nociceptive effects via the µOR would likely occur. With 10 mg/Kg intraperitoneal (i.p) dosing, **56** reached plasma and brain Cmax values of 1690 and 2250 ng/ml and ng/mg—or ∼7 and 10 µM, respectively (**SI Figure 5**). These levels reflect gross exposures, not fraction unbound (Fu). A proxy for the latter is exposure in the cerebrospinal fluid (CSF)^31^ which at 200 nM was about 10-fold over its receptor K_i_, suggesting good coverage. Meanwhile, half-lives for both brain and CSF were between 1.5 and close to 2 hours, suggesting that the compound would persist long enough to support in vivo efficacy experiments.

**Figure 5.**
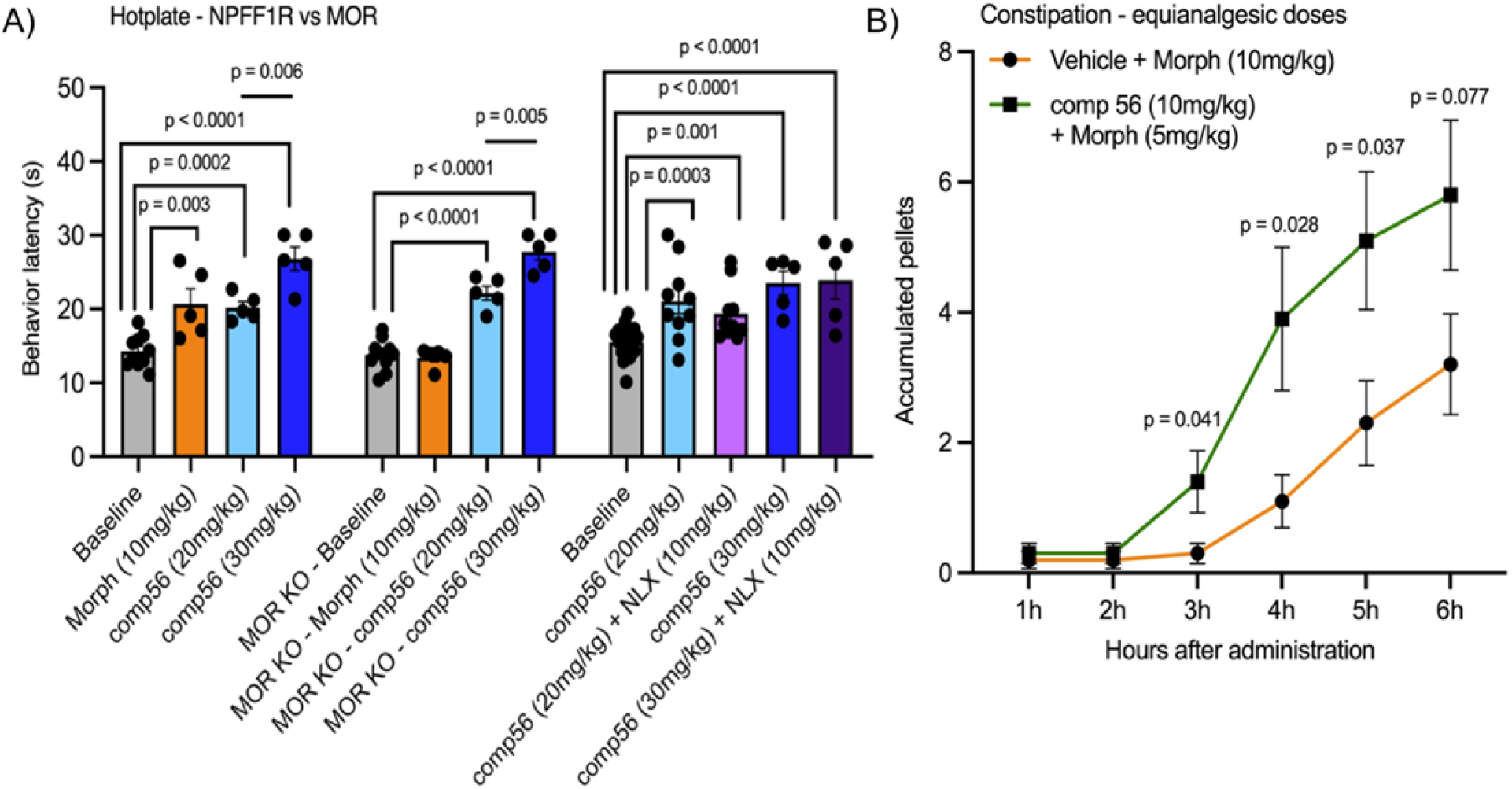
Effects of compound 56 and morphine on analgesia in WT and µOR-knockout mice. (**A**) In the hot-plate assay, the µOR knockout mice experience no analgesia from by i.p. morphine dosing (compare central (10 mg/kg, n=5) and leftmost orange bars (10 mg/kg, n=5), respectively) but do from i.p. dosed compound **56** (compare central light (20 mg/kg, n=5) and dark blue bars (30 mg/kg, n=5) to the same leftmost bars). Furthermore, i.p. naloxone (10 mg/kg) does not inhibit analgesia conferred by i.p. dosed compound **56** at 20 mg/kg (n=10) or 30 mg/kg (n=5, rightmost bars). Data are presented as mean ± SEM. *p*-values were calculated with two-tailed unpaired *t*-tests. (**B**) An equianalgesic dose of i.p. dosed compound **56**+morphine (10 mg/kg + 5 mg/kg, n=10) confers less constipation than i.p. dosed morphine (10 mg/kg, n=10) alone. The effect is visible from three to six hours after administration, and is significant between three and five hours. Data are shown as mean ± SEM. *p*-values were calculated with multiple unpaired *t*-tests at each time point.

### Analgesic activity of compound 56

Compound **56** was dosed (i.p.) at either 10 or 30 mg/Kg in combination with a constant 5 mg/Kg dose of morphine. At this dose, morphine alone induced a modest but significant increase in latency in the tail flick assay, which is thought to represent reflex pain analgesia, over vehicle (**Figure 4A**). Consistent with the hypothesis that an NPFF1R antagonist works to increase the efficacy of µ-opioid agonists^8,9^, addition of 10 and even more so 30 mg/Kg of compound **56** significantly increased the analgesia of this dose of morphine (**Figure 4A**). Meanwhile, compound **56** had no measurable activity on its own in the tail flick assay.

The results in the hot-plate assay, which is thought to measure supraspinally-mediated behavior indicative of a conscious pain experience, were more complicated. As with the tail flick assay, the addition of between 10 and 30 mg/Kg of compound **56** to 5 mg/Kg morphine significantly increased the analgesia in the hot plate assay (**Figure 4B**). Unexpectedly, addition of compound **56** by itself in similar dose ranges, without morphine, also induced strong analgesia (**Figure 4B**), which is inconsistent with the idea that NPFF1R manifests its activities only in cooperation with the µOR.

To investigate the role of the µOR in these effects, we repeated the studies in a µOR knockout mouse (**Figure 5A**). Consistent with cooperative activity between µOR and NPFF1R^8,9^, dosing compound **56** with morphine showed no meaningful analgesia in the tail flick assay in the µOR knockout animal. Conversely, and consistent with some independent action of NPFF1R, in the hot plate assay compound **56** showed little loss in activity in the µOR knockout mouse. Similar results were observed in WT mice when the µOR antagonist naloxone was used to block morphine activity—analgesia was abolished in the tall-flick assay but remained substantial in the hot-plate test (**Figure 5A**)

To control for sedation, which can confound analgesia assays, mice were dosed with **56** with and without morphine and effects on coordination were tested in the rotarod assay (**SI Figure 6**). In most conditions, **56** either alone or in combination with morphine had little effect on mouse coordination. Dosing **56** alone at 30 mg/Kg did lead to a modest decrease in time on the rotarod, consistent with some sedation. However, at lower doses, and in all conditions when dosed with morphine, no significant sedation was observed.

Finally, we asked if the improved analgesia of the **56**/morphine combination would have implications for the adverse drug reactions characteristic of opioid analgesics. Here, also in the mouse, we investigated one of the most common and debilitating side effects of opioid analgesics, namely constipation. At an equianalgesic dose, the combination of **56**+morphine produced less constipation than did morphine dosed alone (**Figure 5B**). This likely reflects the lower morphine doses that are needed to achieve analgesia when combined with a potent NPFF1R antagonist, like compound **56**.

## Discussion

Four results from this study merit emphasis. **First,** from large library docking and structure-based optimization emerged a potent, selective antagonist of the RF-amide receptor NPFF1R, with good physical properties and favorable in vivo exposure. These properties make this molecule suitable as a chemical probe for further biological studies (**Figure 3**). **Second**, the ability to template the docking campaign on a homology model speaks to their ongoing utility in ligand discovery. It also highlights the importance of generating many models and selecting among them based on their ability to prioritize true ligands in control calculations^22,24,25^. **Third,** the selectivity of compound **56** enabled us to use it to probe whether antagonism of NPFF1R will be cooperative with µOR agonism, decreasing nociception, and whether this anti-allodynic effect was strictly cooperative with µOR agonism. While our results support this hypothesis for reflex pain, they also suggest an independent analgesic activity through NPFF1R in the hot-plate assay, which measures cognitively perceived pain (**Figure 4**). In this assay, compound **56** confers analgesia in the absence of morphine, and in both a µOR-knockout and a naloxone-treated mouse (**Figure 5**). **Fourth**, the ability of **56** to complement the activity of morphine enabled lower doses for equianalgesic effects compared to when morphine was used alone. This in turn reduced one of the most common and debilitating side effects of opioid analgesics—constipation—while maintaining strong analgesia (**Figure 5**)

Certain cautions merit airing. Whereas the docking campaign did reveal a 0.3 µM antagonist of NPFF1R, the 4% hit-rate was low versus those against small molecule hormone and neurotransmitter receptors, where hit rates have often been in the 17 to 50% range^15,16,18–21,32^. This likely reflects NPFF1R’s role as a peptide receptor, with its inevitably larger and more shallow binding site, the lack of more than a few drug-like ligands with which to do controls, and errors in our homology model. How these combine to lowering success rates is difficult to quantify, but good comparisons may be to campaigns against the orphan receptors GPR68 and MRGPRX2, which shared several of these features and where hit-rates were also low^24,25^. While it is encouraging that a combination with **56** enables a lower morphine dose, the potential therapeutic impact must be viewed cautiously. Afterall, synergistic combinations of opioids with NSAIDs can still lead to constipation and to addiction.

These caveats should not obscure the key observations of this study. From a docking screen of a diverse, virtual library against a homology model of NPFF1R has emerged a 22 nM antagonist with high selectivity against a large panel of GPCRs. The favorable physical properties of this antagonist contributed to its high brain exposures and to its cooperativity with morphine to improve analgesia. The lower doses of morphine necessary when combined with **56** reduced a key opioid side effect, constipation. The selectivity of **56** for NPFF1R confirms this receptor’s status as a target for the development of antagonists that will act additively or synergistically with opioids, improving analgesia, lowering opioid dose and side-effects.

## Methods

### Homology modeling

Homology models were generated using the RosettaGPCR protocol as previously described^33^ Available crystal structures were analyzed for sequence identity to NPFF1R. Three templates were identified with greater than 30% identity excluding the N-and C-termini: NPY1R (PDB ID: 5ZBQ), OX1R (PDB ID: 4ZJC), OX2R (PDB ID: 4S0V). The sequence of NPFF1R was mapped onto the backbone of the three templates. Further, the coordinates of the co-crystal ligand from NPY1R, UR-MK-299, were truncated to those atoms corresponding to BIBP-3226 and added to each template. The three templates were recombined with each other using a Monte Carlo search algorithm followed by energetic minimization. A set of 1000 models were generated. Models were prepared for docking studies using the blastermaster pipeline^14^ within DOCK3.7^34^.

### Retrospective docking controls

The annotated set of compounds active at NPFF1R were obtained from ZINC20^26 28^. These compounds were sorted by potency and clustered using ECFP4 to obtain a chemically diverse active set. For each active, a set of 50 property-matched decoys were generated from the DUD-E pipeline^26,27^. These decoys are matched for molecular weight, charge, calculated log of the partition between water and octanol (clogP), number of rotatable bonds, and number of hydrogen bond donors and acceptors. The decoys have a scrambled topology with respect to the actives and are therefore presumed to be inactive. A larger set of control ligands were identified by obtaining in-stock compounds within a similar molecular weight and clogP range as the known actives but with no other physical properties controlled for^27^.

### Model selection and optimization

The homology models were challenged to dock and score the diverse active set and property-matched DUD-E decoys. DOCK score was used to rank compounds and models were selected that preferentially ranked actives over decoys. This analysis was quantified by plotting the ranked actives as a function of decoys on a semilogarithmic receiver operator curve. The area under the curve of this semilog plot, or logAUC, was used to rank models. The binding poses of actives for models with high logAUC values were checked visually to ensure they made reasonable interactions within the pocket. Top models were optimized for enhanced discrimination of actives and decoys by altering the parameters of the electrostatic and ligand desolvation scoring grids. Again, top models by logAUC were selected for a small-scale prospective screen on in-stock compounds. A final model was selected that yielded a high number of reasonable hits in the top-scored docking list.

### Large library docking

The ZINC20 database was queried and all monocations with clogP ≥ 1 and a molecular weight > 250 amu were selected to screen at the NPFF1R model. This yielded 166 million protomers for large-scaled docking with DOCK3.7. These were docked on a cluster of 1000 cores. All compounds were rank ordered by DOCK3.7 score, and the top 300,000 were extracted. The compounds were clustered for 2D similarity using ECFP4 chemical fingerprints based on compounds sharing a Tanimoto coefficient (Tc) > 0.35; the single best scoring compound in each cluster was advanced. These were compared to the previously known ligands again using ECFP4 chemical fingerprints and any compound with a Tc > 0.35 to any known was removed. The cluster heads were then re-ranked by DOCK score and the top 5000 were examined visually in the binding pocket. A final set of 26 compounds from among these were selected for experimental testing. These compounds all placed a basic nitrogen within hydrogen bonding distance of Asp^6.59^.

### BRET1 recruitment assay

Using a BRET1 recruitment assay, the agonist and antagonist actions of NPFF1 or NPFF2 drugs were examined. HEK 293T cells were co-transfected with human NPFF1 or NPFF2 with C-terminal Renilla luciferase (RLuc8) and Venus-tagged miniGi at a ratio of 1:4. Transfected cells were plated in plating media (DMEM + 1 % (v/v) dialysed FBS) onto 96-well clear bottom white plates 20–24 hours after transfection. The media was decanted the next day, followed by addition of 75 uL drug buffer (1X HBSS, 20 mM HEPES, 0.1 % (w/v) BSA, pH 7.4) containing a serial dilution of either test ligand or reference compound. After 10 minutes, antagonist assays received 25uL of reference agonist at an EC80 concentration (RFRP-3 for NPFF1 and NPA-NPFF for NPFF2). After another 10 minutes, 25uL of coelenterazine h (Promega) was added to each well to bring the final concentration to 5 uM. Agonist assays are read 20 minutes after media removal; antagonist assays 30 minutes. In a PHERAstar FSX (BMG Labtech), plates were read for luminescence at 485 nm and fluorescence eYFP emission at 530 nm for 1 s per well. The effect of the NPFF1 or NPFF2 drug was represented by calculating the eYFP/RLuc ratio for each well and fitting the net BRET ratio using log(inhibitor) versus response in GraphPad Prism 9.0.

### Tango assay

The PRESTO-Tango assay was used to measure the recruitment of beta arrestin 2 upon ligand stimulation. Tango constructs for NPFF1 and NPFF2 were constructed, and tests were carried out as previously described^35^. Briefly, the NPFF1 or NPFF2 Tango construct was transfected into HTLA cells stably expressing TEV-protease-fused-arrestin (supplied by R. Axel) and a tTA-dependent luciferase reporter gene. The following day, 20,000 transfected cells were put into each well of a 384-well white clear-bottom cell-culture plate coated with poly-l-lysine and filled with DMEM with 1% dialyzed FBS for another 6 hours. For agonist tests, 10 µl of a 5X test compound containing solution was added to each well for an overnight incubation in the same medium as was used for cell plating at a 1X final concentration. For antagonist test, 10 µl of test compound of 1X final concentration is added for 20 mins, additional 10 µl of the EC80 concentration of agonist for NPFF1 or NPFF2 receptor was followed. The next day, wells were filled with 20 µl of Bright-Glo reagent per well after discarding the medium and drug solutions (Promega). After 20 minutes of dark incubation, luminescence was measured for each well on the plate using a Microbeta luminescence reader (Perkin Elmer).

### GPCRome screening

Compounds were screened against the 318 PRESTO-Tango GPCR constructs using previously known techniques, but with a few changes. HTLA cells in DMEM (Sigma) containing 1% (v/v) dialyzed FBS were first plated in white 384-well plates with transparent bottoms. Next day, cells were transfected using PEI (Sigma) utilizing an in-plate modified procedure. In brief, each DNA coding PRESTO-Tango GPCR was resuspended in OptiMEM (Gibco), hybridized with PEI, distributed onto 384-well plates, and then added to cells.

### Animals

Animal experiments were approved by the UCSF Institutional Animal Care and Use Committee and were conducted in accordance with the NIH Guide for the Care and Use of Laboratory animals. Adult (8-10 weeks old) male C56BL/6 mice (strain #664) were purchased from the Jackson Laboratory. Mu-opioid knockout mice were kindly provided by Dr Kevin Yackle at UCSF^32^. Mice were housed in cages on a standard 12:12 hour light/dark cycle with food and water ad libitum.

### Behavioral analyses

The experimenter was always blind to treatment. Compound **56** was dissolved 1h prior to testing in (2-Hydroxypropyl)-β-cyclodextrin (2HPβCD)-saline (20%:80%). Morphine and naloxone were dissolved in saline. For all behavioral tests, animals were first habituated for 30 minutes in Plexiglas cylinders. Mice first received an intraperitoneal (IP) injection of compound **56** followed 30 minutes later by an IP injection of morphine. Behavioral tests were conducted 30 minutes after the morphine injection, as described previously^33^. Briefly, the hindpaw thermal sensitivity was measured by placing the mouse on a 52°C hotplate or, for the tail flick assay, by immersing its tail into a 50°C water bath. For the ambulatory (rotarod) test, mice were first trained on an accelerating rotating rod, 3 times for 5 min, before testing with the compound. All statistical analyses were performed with Prism (Graph Pad).

## Acknowledgments

Supported by Defense Advanced Research Projects Agency grant HR0011-19-2-0020. We thank OpenEye Software for the use of Omega and thank Schrodinger for the use of prepwizard in Maestro.

## Data and compound availability

The structures and identities of compounds docked in this study are freely available from our ZINC20/22 database, http://zinc20.docking.org and https://cartblanche22.docking.org. Raw data are available for all figures. Compound **56** is available from Enamine under registry number Z5075636300.

## Code availability

DOCK3.7 & DOCK3.8 are freely available for non-commercial research http://dock.compbio.ucsf.edu/. A web-based version available to all is available at http://blaster.docking.org/.

