## Supplement Information for "Structure-Based Discovery of a NPFF1R Antagonist with Analgesic Activity"

Table of Contents

Supplemental Figure 1: Model generation and optimization of NPFF1R

Supplemental Figure 2: The 26 compounds selected from virtual screening

Supplemental Figure 3: Aggregation studies of compound **16**

Supplemental Figure 4: Off target screening of compound **56**

Supplemental Figure 5: Pharmacokinetic profile of compound **56**

Supplemental Figure 6: Sedation in the rotarod experiment

Supplemental Table 1: BRET activity of all tested compound **16** analogs

**Supplemental Figure 1: Model generation and optimization of NPFF1R.** (A) The coordinates of BIBP-3226 used in the co-modeling process were obtained from the crystal structure of UR-MK-299 bound to the Neuropeptide Y1 receptor (PDB ID 5ZBQ). (B) The models were biased to have similar orientations of residues Q3.32 and D6.59 as evidenced in the Neuropeptide Y1 crystal structure. (C) There was no correlation between model quality as determined by Rosetta Score and the ability to identify actives from a pool of inactives measured by logAUC. (D) Modification of the dielectric boundary spheres of the top models resulted in varying improvements to the logAUC enrichment.

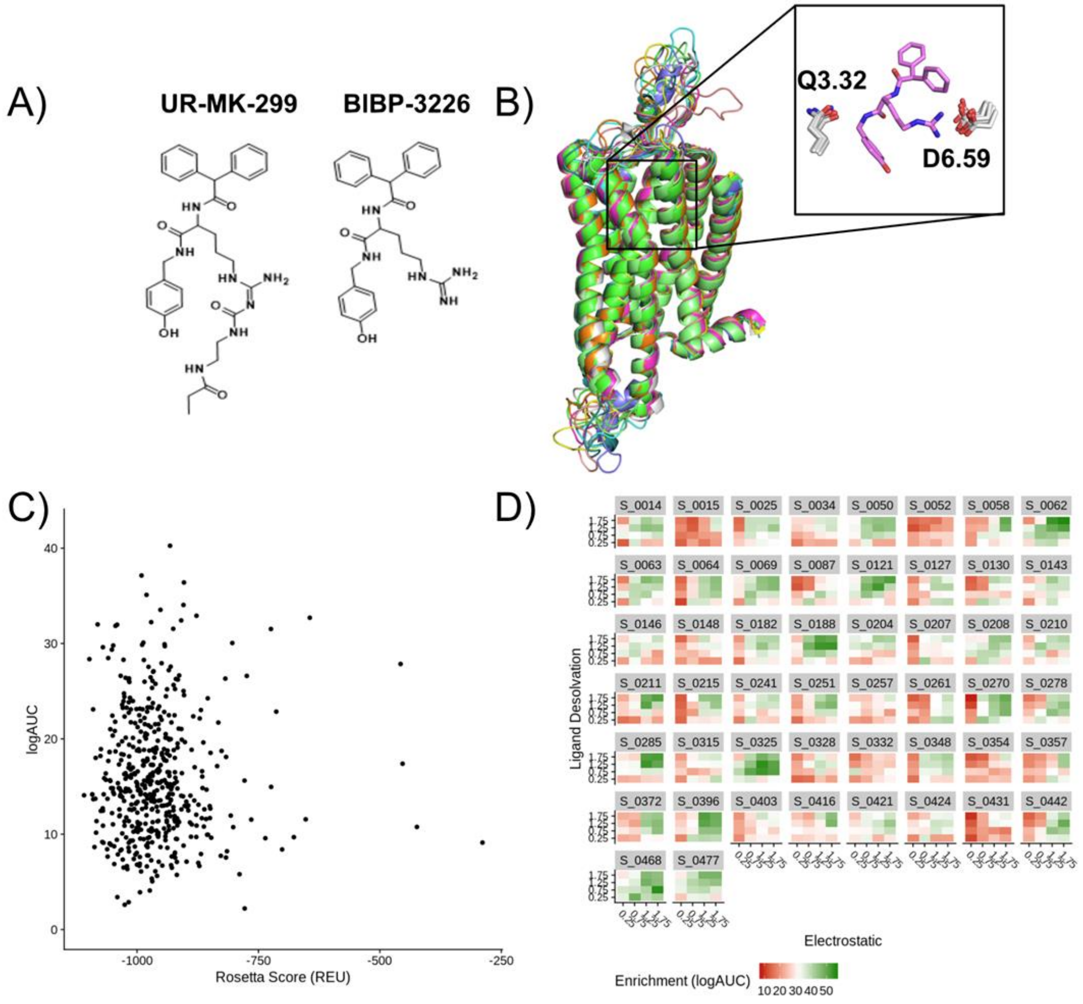

**Supplemental Figure 2: The 26 compounds selected from virtual screening**. The ZINC IDs and 2D structures of all compounds tested from the initial virtual screen.

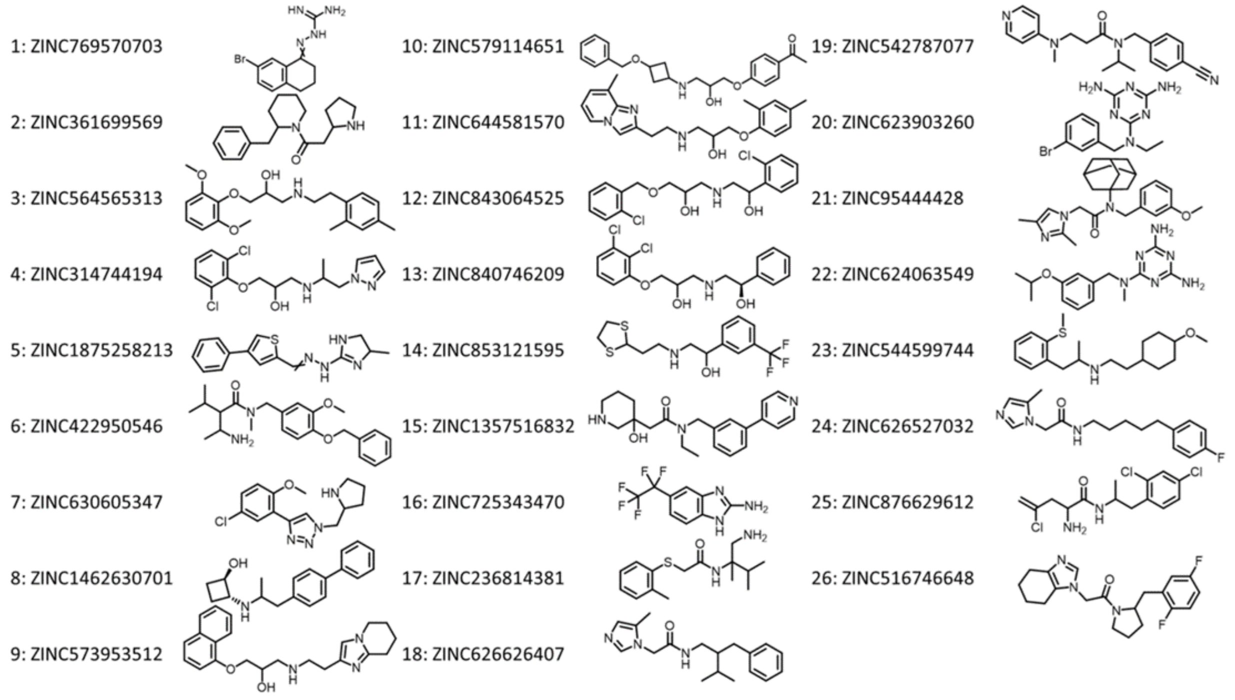

**Supplemental Figure 3: Aggregation studies of compound 16.** (A) Compound **16**, ZINC725343470 or `3470´, was tested in dynamic light scattering for aggregation. The critical aggregation concentration was found to be 12.1 µM, well above the Ki of 319 nM on target activity. (B) Compound **16** was further tested for off-target enzyme inhibition with either MDH or AmpC at 100 µM of ligand. No inhibition was found at either enzyme suggesting the compound activity to be specific for NPFF1R.

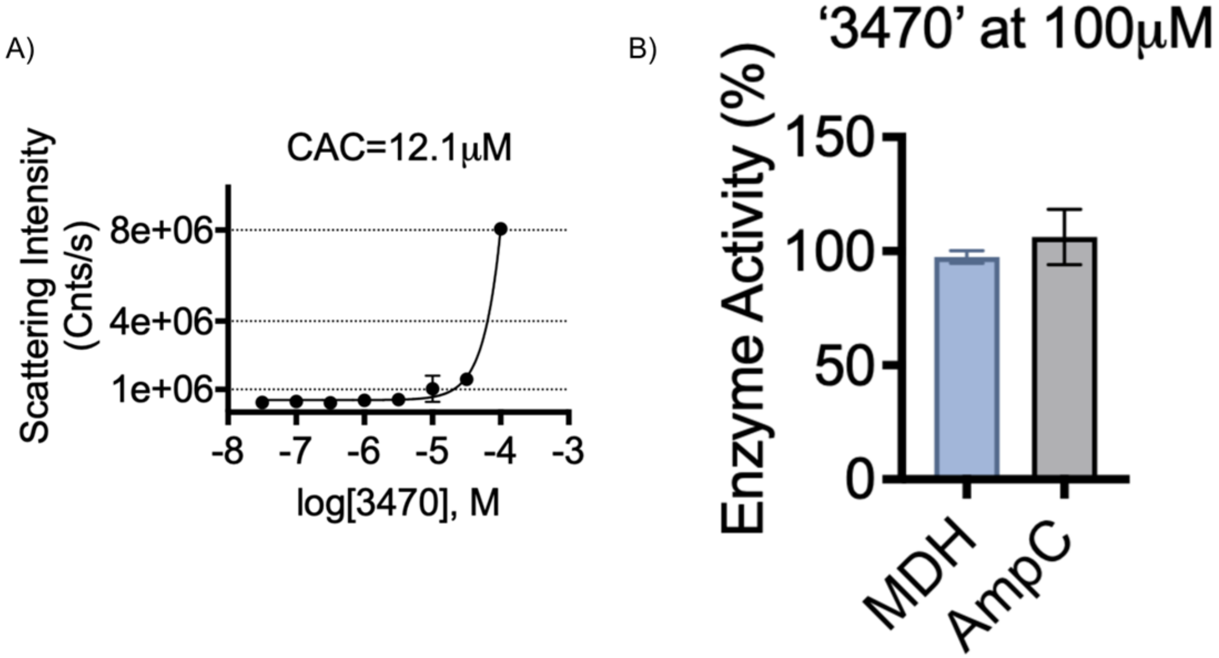

**Supplemental Figure 4: Off target screening of compound 56.** (A) Compound **56** was screened against the GPCRome consisting of 320 GPCRs using a fixed concentration of 3 µM. At this high concentration slight activity was only detected at NPY5R and HRH4. (B) In a concentration response curve, compound **56** displayed weak, non-saturating activity at NPY5R and negligible activity at HRH4. (C) Antagonist activity was measured at NPY1R for compound **56** and other antagonists. BIBP-3226 shows measurable activity at NPY1R demonstrating its non-selectivity at this target. The remaining compounds, including compound **56**, demonstrate relatively weak activity.

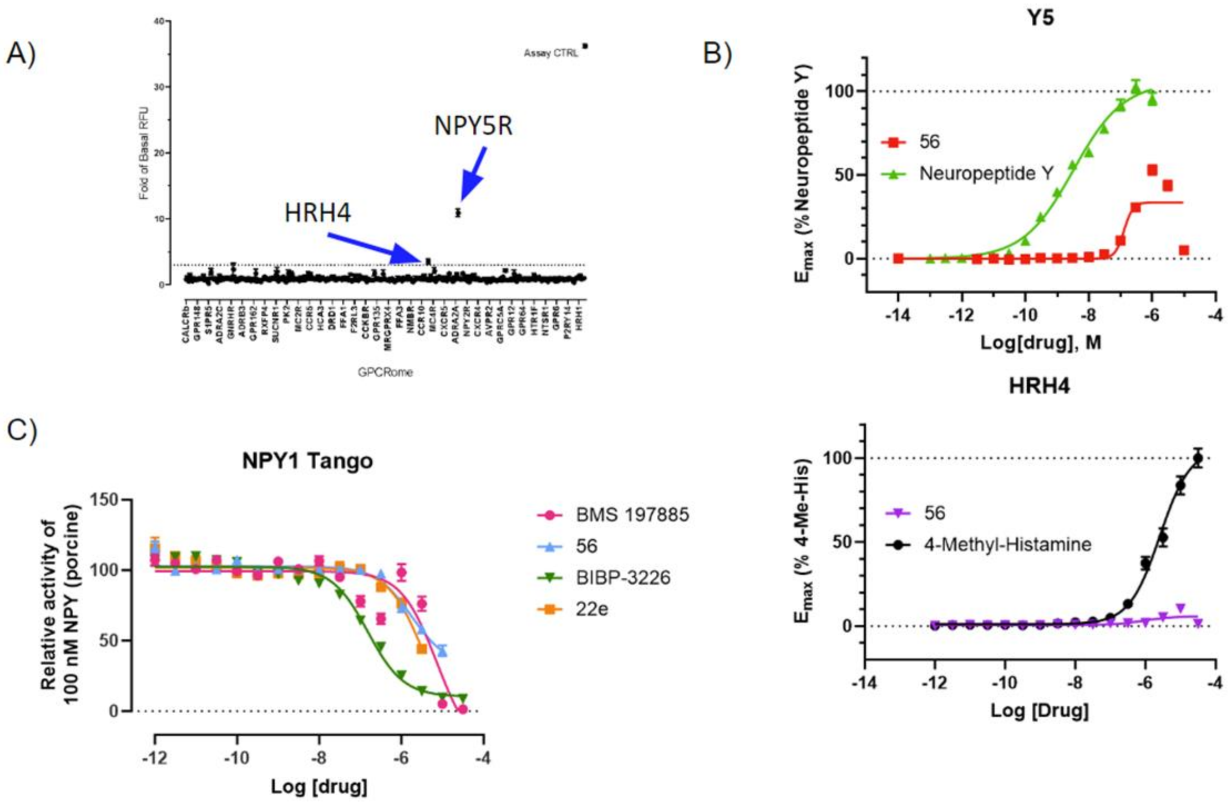

**Supplemental Figure 5: Pharmacokinetic profile of compound 56.** Following IP injection of 10mg/kg compound 56, time course measurements were taken showing an accumulation of the compound in brain after 2 hours.

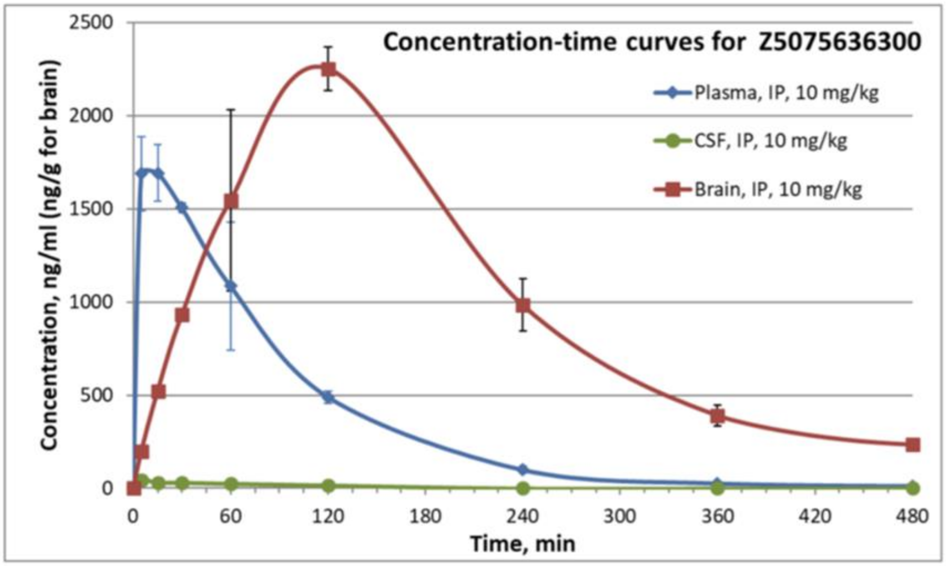


| Sample | Administration | Dose, mg/kg | Pharmacokinetic Parameters | | | | | |
|---|---|---|---|---|---|---|---|---|
| | | | Tmax, Min | Cmax, ng/ml (g) | AUC$_{0\to t}$ (AUClast) ng*min/ml (g) | AUC$_{0\to\infty}$ (AUCINF_obs), ng*min/ml (g) | $T_{1/2}$ (HL_Lambda_z), min | $K_{el}$ (Lambda_z), min$^{-1}$ |
| Plasma | IP | 10 | 15.0 | 1690 | 176000 | 177000 | 61.6 | 0.0112 |
| Brain | IP | 10 | 120 | 2250 | 480000 | 517000 | 108 | 0.00642 |
| CSF | IP | 10 | 5.00 | 46.0 | 3000 | 5230 | 97.4 | 0.00712 |

**Supplemental Figure 6: Sedation in the rotarod experiment.** Compound **56** confers no apparent sedation up to 30 mg/Kg in isolation. No measurable sedation was observed in combination with morphine.

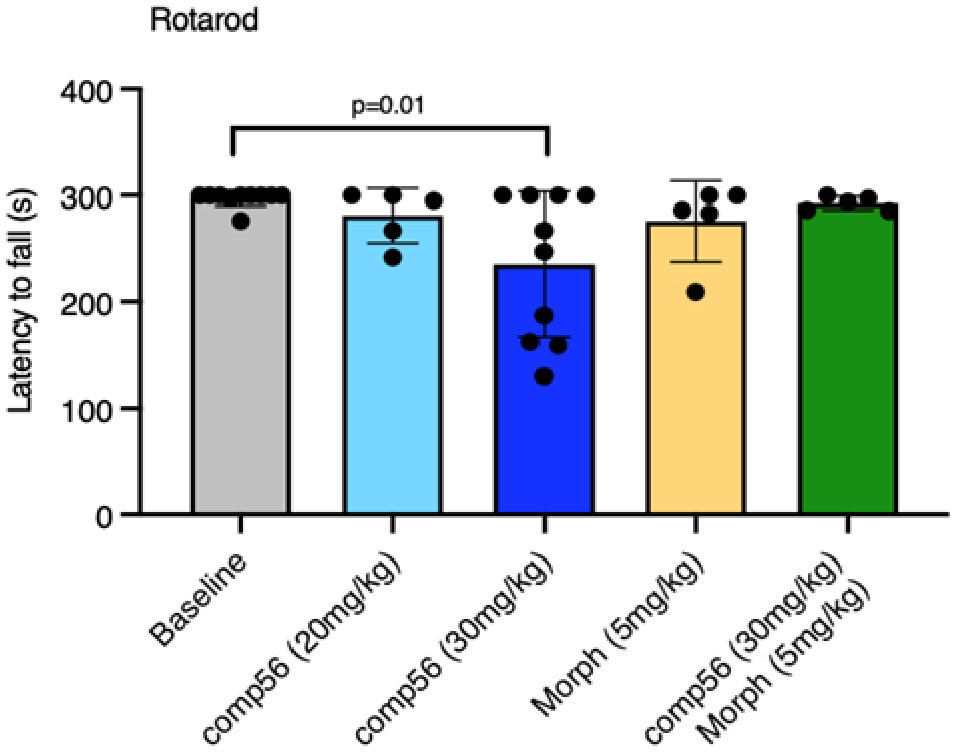

Supplemental Table 1: BRET activity of all tested compound 16 analogs.

| Compound ID | 2D Structure | Ki (nM) | Enamine ID* | SMILES |
|---|---|---|---|---|
| **16** | 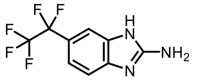 | 319 | Z3016357374 | NC1=NC=2C=C(C=CC2N1)C(F)(F)C(F)(F)F |
| **27** | 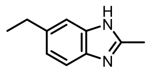 | inactive | Z3241580504 | CCC=1C=CC=2N=C(C)NC2C1 |
| **28** | 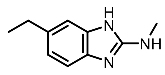 | inactive | Z2751251823 | CCC=1C=CC=2N=C(NC)NC2C1 |
| **29** | 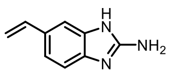 | 650 | Z4420693782 | NC1=NC=2C=CC(C=C)=CC2N1 |
| **30** | 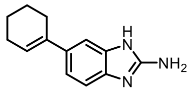 | 342 | Z4420693967 | NC1=NC=2C=CC(=CC2N1)C3=CCCCC3 |
| **31** | 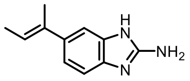 | 123 | Z4416333218 | C/C=C(\C)/C=1C=CC=2N=C(N)NC2C1 |
| **32** | 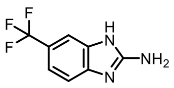 | inactive | Z1511494445 | NC1=NC=2C=C(C=CC2N1)C(F)(F)F |
| **33** | 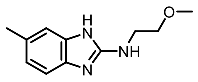 | inactive | Z56807718 | COCCNC1=NC=2C=C(C)C=CC2N1 |
| **34** | 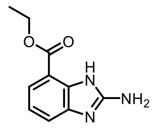 | inactive | Z1509585071 | CCOC(=O)C=1C=CC=C2NC(N)=NC12 |
| **35** | 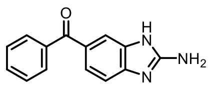 | inactive | 35403 (Sigma) | C1=CC=C(C=C1)C(=O)C2=CC3=C(C=C2)N=C(N3)N |
| **36** | 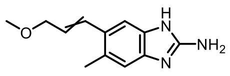 | inactive | Z4223307867 | COCC=CC=1C=C2NC(N)=NC2=CC1C |
| **37** | 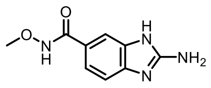 | inactive | Z4874003063 | CONC(=O)C=1C=CC=2NC(N)=NC2C1 |
| **38** | 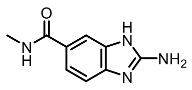 | inactive | Z4874003067 | CNC(=O)C=1C=CC=2NC(N)=NC2C1 |
| **39** | 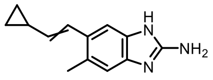 | inactive | Z4223832984 | CC=1C=C2N=C(N)NC2=CC1C=CC3CC3 |
| **40** | 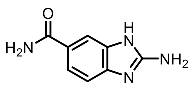 | inactive | Z1262646398 | NC(=O)C=1C=CC=2NC(N)=NC2C1 |
| **41** | 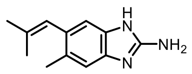 | 178 | Z4223213409 | CC(=CC=1C=C2NC(N)=NC2=CC1C)C |
| **42** | 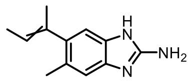 | 25 | Z4224145264 | CC=C(C)C=1C=C2NC(N)=NC2=CC1C |

| Compound ID | 2D Structure | Ki (nM) | Enamine ID* | SMILES |
|---|---|---|---|---|
| **43** | 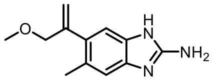 | 197 | Z4223596179 | COCC(=C)C=1C=C2NC(N)=NC2=CC1C |
| **44** | 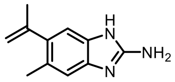 | 48 | Z4874090623 | CC(=C)C=1C=C2NC(N)=NC2=CC1C |
| **45** | 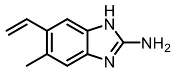 | 255 | Z4874088327 | CC=1C=C2N=C(N)NC2=CC1C=C |
| **46** | 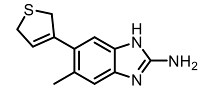 | 133 | Z4223455428 | CC=1C=C2N=C(N)NC2=CC1C3=CCSC3 |
| **47** | 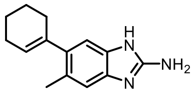 | 664 | Z4874003068 | CC=1C=C2N=C(N)NC2=CC1C3=CCCCC3 |
| **48** | 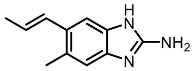 | 222 | Z4874088337 | C/C=C/C=1C=C2NC(N)=NC2=CC1C |
| **49** | 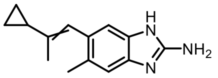 | 260 | Z4874003071 | CC(=CC=1C=C2NC(N)=NC2=CC1C)C3CC3 |
| **50** | 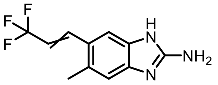 | 2655 | Z4874003072 | CC=1C=C2N=C(N)NC2=CC1C=CC(F)(F)F |
| **51** | 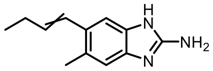 | 588 | Z4874003073 | CCC=CC=1C=C2NC(N)=NC2=CC1C |
| **52** | 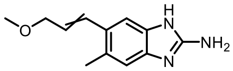 | 4671 | Z4223307867 | COCC=CC=1C=C2NC(N)=NC2=CC1C |
| **53** | 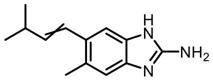 | 1170 | Z4874003074 | CC(C)C=CC=1C=C2NC(N)=NC2=CC1C |
| **54** | 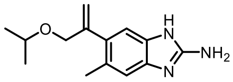 | 75 | Z4342120597 | CC(C)OCC(=C)C=1C=C2NC(N)=NC2=CC1C |
| **55** | 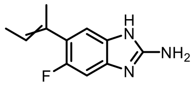 | 1705 | Z5064372854 | CC=C(C)C=1C=C2N=C(N)NC2=CC1F |
| **56** | 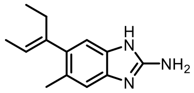 | 22 | Z5075636300 | CC/C(=C\C)/C=1C=C2NC(N)=NC2=CC1C |
| **57** | 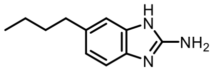 | 50 | Z1216815549 | CCCCC=1C=CC=2NC(N)=NC2C1 |
| **58** | 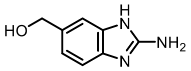 | inactive | Z1263714733 | NC1=NC=2C=C(CO)C=CC2N1 |
| **59** | 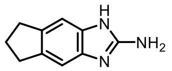 | 70 | Z5340651745 | NC1=NC=2C=C3CCCC3=CC2N1 |

* Enamine ID unless otherwise noted
